# Lung *Mycobacterium tuberculosis* infection perturbs metabolic pathways in non-pulmonary tissues

**DOI:** 10.1101/2024.02.09.579656

**Authors:** Falak Pahwa, Shweta Chaudhary, Anushka Dayal, Ranjan Kumar Nanda

**Affiliations:** Translational Health Group, International Centre for Genetic Engineering and Biotechnology, New Delhi-110067, India; TERI School of Advanced Studies, New Delhi-110070, India

**Keywords:** Tissue metabolites, GC-MS, TB infection, Fecal metabolomics, Amino acid metabolism

## Abstract

*Mycobacterium tuberculosis* (Mtb), through aerosol, reaches the lungs to cause pulmonary tuberculosis (TB); however, it may also affect the metabolism of other tissues in age-specific ways. In this study, female C57BL/6 mice (2 and 5 months old; M) were infected with a low aerosol dose (100-200 cfu) of Mtb H37Rv to monitor tissue mycobacterial load and multi-tissue metabolite profiling using gas chromatography and mass spectrometry (GC-MS). 5M C57BL/6 mice showed separate tissue metabolic phenotype with significantly higher lung aspartic acid, fecal oxalic acid and tryptophan levels with lower liver lysine and aspartic acid and fecal phenylalanine levels (log_2_FC: 5M/2M> ±1.0, p<0.1) compared to 2M young controls. Upon Mtb infection, the lung mycobacterial load of 2M and 5M mice were similar till 6 weeks post-infection. However, significantly higher lung phosphoric acid, malonic acid and lower mannose levels (log_2_FC: Mtb infected/healthy> ±1.0, p<0.1) were observed in Mtb-infected 5M C57BL/6 mice. Meanwhile, Mtb-infected 2M mice showed higher liver xylose and lower lysine levels. The thigh muscles of Mtb-infected 2M and 5M mice showed increased malic acid and oxalic acid and decreased glycine, serine, and glycerol levels. Fecal aspartic acid level was higher in Mtb-infected 5M mice, while a decreased abundance of fecal lysine was observed in Mtb-infected 2M mice. Overall, this study demonstrates a deregulated tissue-specific amino acid metabolism in Mtb-infected mice groups of different age groups, which might be targeted for managing TB infection-related adverse effects.

## Introduction

*Mycobacterium tuberculosis* (Mtb) infection causes tuberculosis (TB) disease and contributes to a total of 1.78 million new young (10-24 years) patients annually.^1^ TB incidence is reported to increase rapidly during adolescence, with a peak in early adulthood (20-24 years) in high TB endemic regions.^1,2^ To understand TB-related pathophysiology, C57BL/6, BALB/c and C3HeB/FeJ mice strains of 6-8 weeks of age representing ∼18 years of human age are commonly used.^3^ Host age plays a critical role in controlling the pathogen, and the effect of the pathogen may not necessarily be limited to the specific organ. Thus, understanding disease manifestation and its impact in multiple tissues of different age groups of model animals is critical.

Several reports on biofluid metabolomics have shown metabolic dysregulation in Mtb-infected mice or humans. Tissue-specific metabolome changes were reported in the lungs of the Mtb-infected C57BL/6 mice model.^4,5^ However, a detailed characterization of the amino acid profiles in other host tissues of Mtb-infected mice in different age groups is missing in the literature. Identifying perturbed pathways in the host tissues that do not directly encounter the mycobacteria might be useful in identifying targets for benefitting TB patients.

In this study, female C57BL/6 mice of 2 and 5 months old were aerosol infected with Mtb H37Rv strain and tissue mycobacterial load was monitored. Along with lung, liver, thigh muscle, brain and fecal metabolites were profiled using gas chromatography-mass spectrometry (GC-MS). Overall, our data demonstrates that post-Mtb infection amino acid metabolism is deregulated, which is more pronounced in 5M mice. Targeting amino acid metabolic pathways will be useful in developing adjunct therapeutics for TB.

## Materials and Methods

### Experimental groups and Mtb infection

Procedures adopted in this study were performed following the recommended Guidelines for Care and Use of Laboratory Animals and approved by the Animal Ethics Committee of the International Centre for Genetic Engineering and Technology (ICGEB), New Delhi (Ref No. ICGEB/IAEC/07032020/TH-10). Female C57BL/6 mice of two age groups (2 and 5 months: M) were procured from the ICGEB bio-experimentation facility and transferred to the Tuberculosis Aerosol Challenge Facility (TACF; BSL-3 facility) for infection experiments. Throughout the experiments, mice were fed a chow diet and water ad libitum and maintained at 20-22 °C, 45-60% humidity with a 12 h light/dark cycle. Weight-adjusted randomly grouped mice (n = 4 per group), after 1 week of acclimatization in TACF, were infected with a low dose (100-120 cfu) of animal passaged *Mycobacterium tuberculosis* H37Rv strain in Madison aerosol exposure chamber (University of Wisconsin, USA). Mice were euthanized by isoflurane overdose followed by cervical dislocation under the supervision of a veterinarian, and a necropsy was carried out. Tissues (lung, spleen and liver) were harvested for bacterial cfu assay at indicated time points. A part of the fresh tissue (lung, liver, thigh muscle and brain) was treated with chilled methanol (kept at -80 °C) for metabolite analysis and stored at -80 °C. Fresh fecal pellets of mice belonging to different study groups were collected (two times a day: morning and evening) by holding the mice by the tail base on disinfected aluminium foil and collected in a tube at indicated time points and stored at -80 °C until further use.

### Bacterial load monitoring by cfu assay

Briefly, part of the tissues (left lobe of lungs, the upper half of spleen, and two-thirds of liver) were homogenized using T-25 digital ultra-turrax disperser (IKA, India) in phosphate-buffered saline (PBS, 1 mL). The tissue homogenates were serially diluted and plated in duplicates on 7H11 agar supplemented with Middlebrook OADC (oleic acid, albumin, dextrose and catalase, 10%, BD Biosciences, #211886). These plates were incubated at 37 ºC in a humidified incubator, and bacterial colonies were enumerated at 3 weeks post-inoculation.

### Tissue metabolite analysis using mass spectrometry

Stored tissue samples (∼100 mg from lungs, liver, thigh muscle and brain) were taken in a 2 mL bead beating tube containing zirconium beads (250 mg, 2 mm diameter, BioSpec Products, #11079124zx). After adding ribitol (2 mg/mL, 5 μL), chilled methanol (80%, 1 mL) was added. Bead beating (6 cycles, 1 min on/off, on ice between cycles, 6500 Hz speed/cycle) was performed, and the solution was kept on ice for 30 min. The extracted tissue metabolites were centrifuged at 10,000 × g at 4 °C for 10 min. The supernatant was passed through a filter (0.2 μm; 25 mm diameter Nylon for infected lung, liver and brain; 4 mm diameter PVDF for infected thigh muscle). A quality control (QC) sample was prepared by pooling equal volumes (∼110 μL) of the extracted tissue metabolites from all the samples (n=64). A part of the methanolic fraction (∼400 μL) was vacuum dried using SpeedVac (Labconco, USA) at 40 °C until complete dryness. Methoximation of these dried metabolites were carried out with methoxyamine-hydrochloric acid (MOX-HCl, 2%, 40 μL, Supelco, #33045-U) and incubated for 2 h at 60 °C at 400 rpm on a thermomixer (Eppendorf ThermoMixer C, Germany, #EP5382000023). Then, derivatization was done with N-Methyl-N-(trimethylsilyl) trifluoroacetamide (MSTFA, 70 μL, Supelco, #69479) and incubated for 30 min at 60 °C at 400 rpm on thermomixer. The mixture was again centrifuged at 16,000 × g at 4 °C for 10 min. The supernatant was transferred to a glass vial insert (200 μL) in a GC vial (2 mL capacity) for mass spectrometry data acquisition using a Gas Chromatography-Time of flight-Mass Spectrometry (GC-TOF-MS, LECO Pegasus, USA). Derivatized metabolites (1 μL) were injected in a splitless mode using helium as carrier gas with a flow rate of 1 mL/min into RTX-5 column (30 m × 0.25 mm × 0.25 μm; Restek, USA). Metabolite separation was carried out using a temperature gradient of 50 °C for 1 min, followed by a ramp of 8.5 °C/min to 200 °C and further a ramp of 6 °C/min till 280 °C and hold time for 5 min. Mass spectrometric data acquisition was carried out at −70 eV, and a range of m/z 45 to 600 was scanned at 20 spectra/s. A solvent delay of 600 s was used before GC-MS data acquisition. Detector voltage and ion source temperature were set at 1678 V and 225 °C respectively. All GC-TOF-MS parameters were controlled by ChromaTOF software (version 4.50.8.0; Leco, USA), and all sample data acquisition was completed within 24 h of derivatization. Commercial standards of important molecules were processed using similar methods used for sample analysis to confirm the identity of important amino acids.

Acquired data files of tissue samples were aligned using “Statistical Compare” feature of ChromaTOF. A retention time tolerance of 0.5 s, minimum spectral similarity of 600 and the signal-to-noise ratio (S/N) threshold was set as 75 to align the molecular features. Features qualifying set of parameters and their abundance were selected as metadata, and manual data curation was carried out for peak alignment. The selection criteria comprised of removing the analytes with signal: noise<75, reagent peaks, low baseline analytes, and low library matches (<600). Analytes showing the same mass fingerprints eluting within a retention time of 20 seconds were merged as a single analyte. Library from the National Institute of Standards and Technology, NIST (version 11.0) consisting of mainlib (212,961 spectra) and replib (30,932 spectra) was used for tentative molecular feature identification. After manual data matrix curation from ∼409 identified analytes, 99 were considered, data normalized using internal standard ribitol, followed by statistical analysis using MetaboAnalyst (version 5.0).

### Fecal metabolite analysis

Fecal pellets (∼50 mg) were weighed, taken in a bead beating tube containing zirconia beads (300 μL, 0.1 mm diameter, BioSpec Products, #11079101z) and to it, chilled methanol (80%, 1 mL, kept at -20 °C) was added along with ribitol (2 mg/mL, 5 μL). Bead beating (3 cycles, 30 s on/off, on ice between cycles, 6500 Hz speed/cycle) was carried out, and the mixture was centrifuged at 16,000 × g at 4 °C for 15 min. The supernatant was passed through a nylon filter (0.2 μm) and transferred to a fresh tube, and a QC sample was prepared by pooling equal volumes (100 μL) of all the extracted fecal metabolites (n=25). All the extracted fecal metabolites were vacuum-dried using SpeedVac at 40 °C until complete dryness. Methoximation of these dried metabolites was carried out with methoxyamine-hydrochloric acid (MOX-HCl, 3%, 20 μL, Supelco, #33045-U) and incubated for 90 min at 37 °C at 1000 rpm on a thermomixer. Then, derivatization was done with N-Methyl-N-(trimethylsilyl) trifluoroacetamide (MSTFA) and trimethylchlorosilane (1% TMCS, 80 μL, Supelco, #69478) and incubated for 30 min at 37 °C at 1000 rpm on thermomixer. The mixture was again centrifuged at 10,000 × g at 4 °C for 5 min. The supernatant was transferred to a glass vial insert (200 μL) in a GC vial (2 mL capacity) for GC-MS data acquisition.

Derivatized fecal metabolites (1 μL) were injected in a splitless mode using helium as carrier gas with a flow rate of 0.5 mL/min to RTX-5 column (30 m × 0.25 mm × 0.25 μm; Restek, USA). The temperature parameters: 60 °C for 1 min, followed by a ramp of 10 °C/min to 220 °C and further a ramp of 5 °C/min till 300 °C and a hold time for 5 min were used for metabolite separation. Mass spectrometric data acquisition was carried out at −70 eV, and a mass range of m/z 50 to 600 was scanned in normal mode with a gain factor of 1 with a solvent delay of 6.8 min. Detector voltage and ion source temperature were set at 1388 V and 230 °C respectively. All GC-MS parameters were controlled by ChemStation (G1701EA E.02.02.1431; Agilent Technologies), and data acquisition was completed within 24 h of derivatization. Commercial standards were processed using a similar method to confirm the identity of the important amino acid analytes.

Acquired data files of fecal samples were converted from .d to .elu format using AMDIS (Automated Mass Spectral Deconvolution and Identification System). All .elu files were uploaded into the SpectConnect web server software for data matrix preparation^12^. Using SpectConnect, we obtained 365 features and focused on 20 analytes, whose identity was confirmed by running commercial standards (**Supplementary Figure S1**). The resulting matrix included the retention time of the representative peak for each metabolite, and the peaks were tentatively identified by searching against the National Institute of Standards and Technology, NIST (version 11.0) database consisting of mainlib (243,893 spectra) and replib (30,932 spectra). MetaboAnalyst (version 5.0) was used for multivariate analysis using the metadata to identify deregulated metabolic features.

### Statistical Analysis

The body weight and CFU data were analyzed by parametric analysis using Student’s t-test in GraphPad Prism (version 8.4.2, GraphPad Software, San Diego, CA). All data are presented as mean ± standard deviation (S.D.). A p-value of ≤0.05 was considered significant at a 95% confidence interval. Information about the sample size and statistical analysis adopted in each experiment has been provided in the individual figure legend.

## Results

### Optimization of GC-MS parameters and sample volume for tissue metabolite analysis

In an optimization experiment using various tissues (lungs, spleen, liver, brain, heart, thigh muscle, and kidneys) of healthy 2M female C57BL/6 mice, the total ion chromatogram (TIC) from the extracted polar metabolites from these tissues showed differences in their profile (**Supplementary Figure S2A**). Partial least squares-discriminant analysis (PLS-DA) plots of these tissues showed separate clusters and, thus, a tissue-specific metabolome (**Supplementary Figure S2B**). We observed high proline levels in the liver, which is essential for hepatocyte growth and higher glutamine in the brain, which is a precursor to the neurotransmitter glutamate (*data not shown*). Thigh muscle had higher levels of isoleucine, which regulates body weight and muscle protein synthesis (*data not shown*). Thus, we asked whether tissue-specific metabolome changes upon Mtb infection.

QC of metabolites extracted from tissues of H37Rv Mtb-infected 2M and 5M female C57BL/6 mice and further derivatized and acquired on GC-TOF-MS (**Figure 1A**). Amongst different QC volumes (25, 50, 75, 100, 200, 400 and 800 μL) used, 400 μL yielded optimum results (**Figures 1B** and **C**). Similarly, 400 μL metabolites extracted from the lungs, liver, thigh muscle and brain yielded optimum metabolic features (**Supplementary Figure S3**). Employing the optimized GC-MS method parameters, an injection of 1 μL of QC sample volume yielded optimum information (**Figure 1D**).

**Figure 1.**
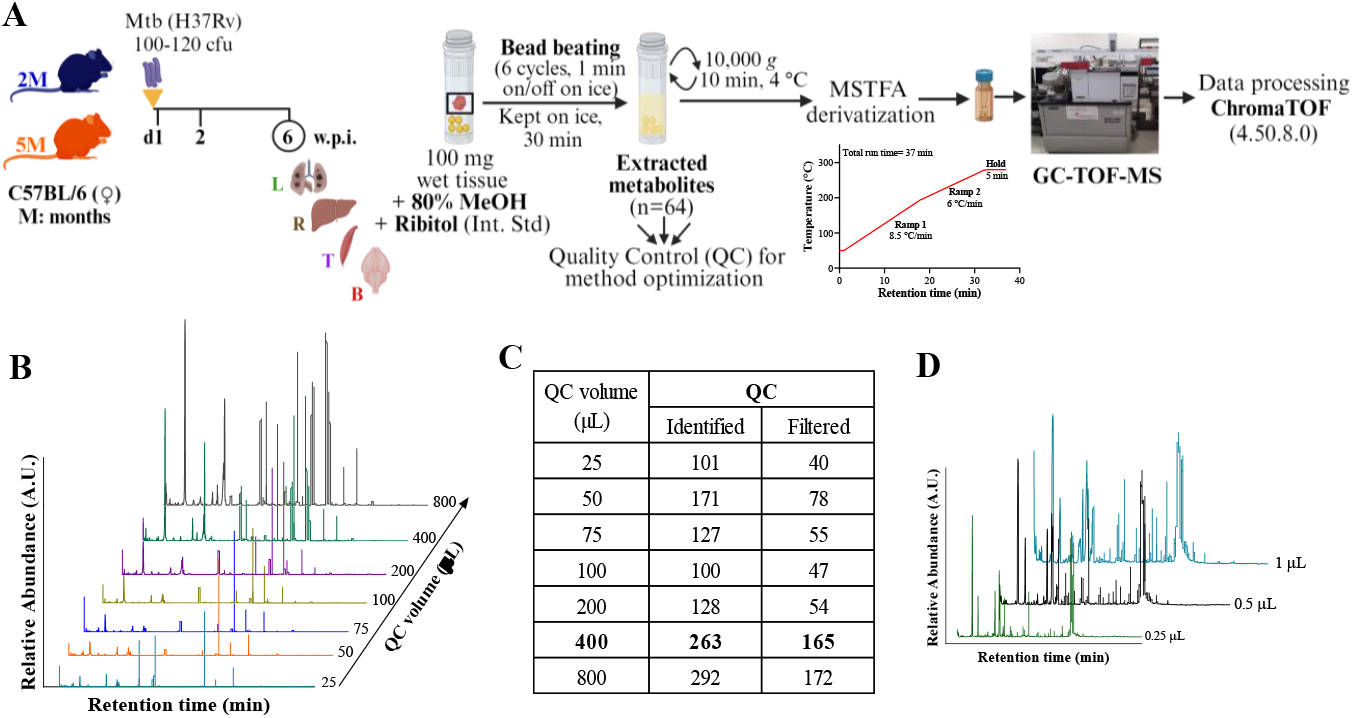
Optimization experiments for tissue metabolomics. **A**. Schematic representation of tissues (L: Lungs, R: Liver, T: Thigh muscle, B: Brain) taken for metabolite profiling and the protocol used for tissue metabolomics experiment: Extraction of tissue metabolites, processing of metabolites for Gas Chromatography-time of flight-mass spectrometry (GC-TOF-MS). **B**. Total ion chromatograms (TIC) of different volumes of quality control (QC) used. **C**. Table showing the identified and filtered analytes of different volumes of QC samples. **D**. TIC of samples using different injection volumes. M: months; 2M in blue and 5M in orange; AU: arbitrary units, min: minutes.

### Young (2M/5M, female) C57BL/6 mice showed minor changes in bacterial burden upon Mtb infection

Lung mycobacterial burden of Mtb-infected 2M and 5M mice showed a time-dependent increase from 2 to 6 w.p.i. (**Figure 2A**). However, the lung Mtb burden was comparable between the age groups at 2 and 6 w.p.i. (**Figure 2A**). Spleen and liver of 2M and 5M mice showed similar Mtb load at both time points (**Figure 2A**). Since the lung Mtb load was similar, we hypothesized metabolic dysregulation upon Mtb infection in multiple tissues of both age groups.

**Figure 2.**
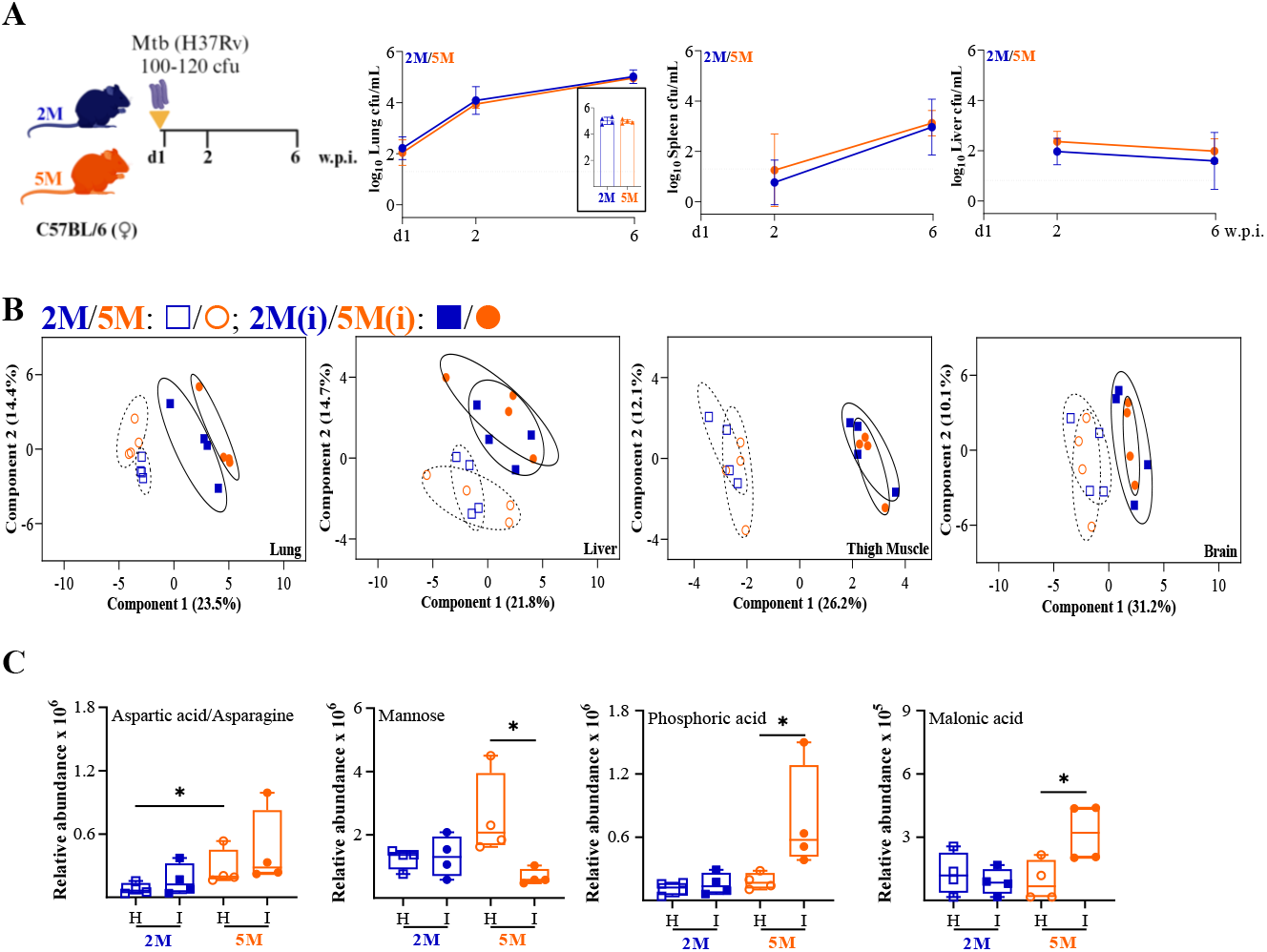
Deregulation in lung tissue metabolome of Mtb infected 5M C57BL/6 mice. **A**. Schematic of the experimental details used in this experiment including C57BL/6 mice (female; age groups of 2 and 5 months; M). Bacterial burden in the lung, spleen and liver of Mtb infected mice at 2 and 6 w.p.i. showing mean of n=4/time point/age group, with error bars (standard deviation); dashed horizontal line represents limit of detection (LOD); Individual data points of lung cfu at 6 w.p.i. are presented as histogram in the inset. **B**. Partial least squares-discriminant analysis (PLS-DA) plot representing tissue (lung, liver, thigh muscle and brain) specific metabolic differences in Mtb infected (I, solid symbols) and healthy (H, open symbols) C57BL/6 mice of two age groups. **C**. Relative abundances of lung metabolites presented as histograms. n = 4/tissue/age group/condition; M: months; 2M in blue and 5M in orange; w.p.i.: weeks post infection; p-value: * ≤0.01 at 90% confidence interval.

### Aerosol Mtb infection leads to deregulation in liver and thigh muscle of Mtb-infected mice groups

QC samples clustered together in the principal component analysis (PCA) plot indicating minimal method-associated variation (**Supplementary Figure S4A**). The tissue-specific metabolite matrices were prepared for both age groups, and the number of variables used for analysis is presented in **Supplementary Table S1**. The metabolites extracted from the liver, thigh muscle and brain of healthy mice of both age groups clustered together (**Figure 2B, Supplementary Figure S4A**). However, aspartic acid/asparagine was found to be higher (log_2_FC: Mtb infected/healthy=1.2) in the lungs of 5M healthy mice (**Figure 2C**). Meanwhile, the livers of 5M mice had a lower abundance of lysine (log_2_FC= -1.1) and aspartic acid/asparagine (log_2_FC= -1, **Supplementary Figure S5A**). Upon Mtb infection, lung and liver metabolites of mice showed age-specific clusters in the PLS-DA plot (**Figure 2B, Supplementary Figure S4B**). While, thigh muscle and brain metabolites of Mtb-infected 2M and 5M mice had overlapping clusters (**Figure 2B, Supplementary Figure S4B**). The abundance of phosphoric acid (log_2_FC=1.8) and malonic acid (log_2_FC=1.5) were found to be higher, while mannose (log_2_FC= -2.1) was lower in the lungs of Mtb-infected 5M mice compared to healthy controls (**Figure 2C**). The liver of Mtb-infected 2M mice showed higher xylose (log_2_FC=2) and lower lysine (log_2_FC= -1) compared to healthy mice (**Supplementary Figure S5A**). Significant loss of glycine, serine and glycerol was observed in the thigh muscle of both age groups upon Mtb infection (**Supplementary Figure S5B**). In contrast, malic acid and oxalic acid were increased in the thigh muscle of Mtb-infected mice of both age groups (**Supplementary Figure S5B**). Interestingly, brain metabolites of Mtb-infected mice, irrespective of the age groups, showed minimal variations (**Figure 2B**).

### Altered fecal metabolome in 2M C57BL/6 mice upon Mtb infection

We next asked how fecal metabolome is altered upon healthy ageing and Mtb infection (**Figure 3A**). PLS-DA plot showed close clustering of the QC samples, while fecal metabolites of healthy 2M and 5M and Mtb-infected groups showed separate clusters (**Supplementary Figure S6**). Importantly, oxalic acid (log_2_FC=2.9) and tryptophan (log_2_FC=1.8) showed increased abundance, while phenylalanine (log_2_FC= -1) decreased with aging (**Figure 3B, Supplementary Figure S7**). Amino acids, including lysine (log_2_FC= -1.1) were decreased in Mtb-infected 2M mice, while aspartic acid (log_2_FC=1) levels were increased in the fecal samples of Mtb-infected 5M mice compared to healthy controls (**Figures 3C** and **D, Supplementary Figure S7**).

**Figure 3.**
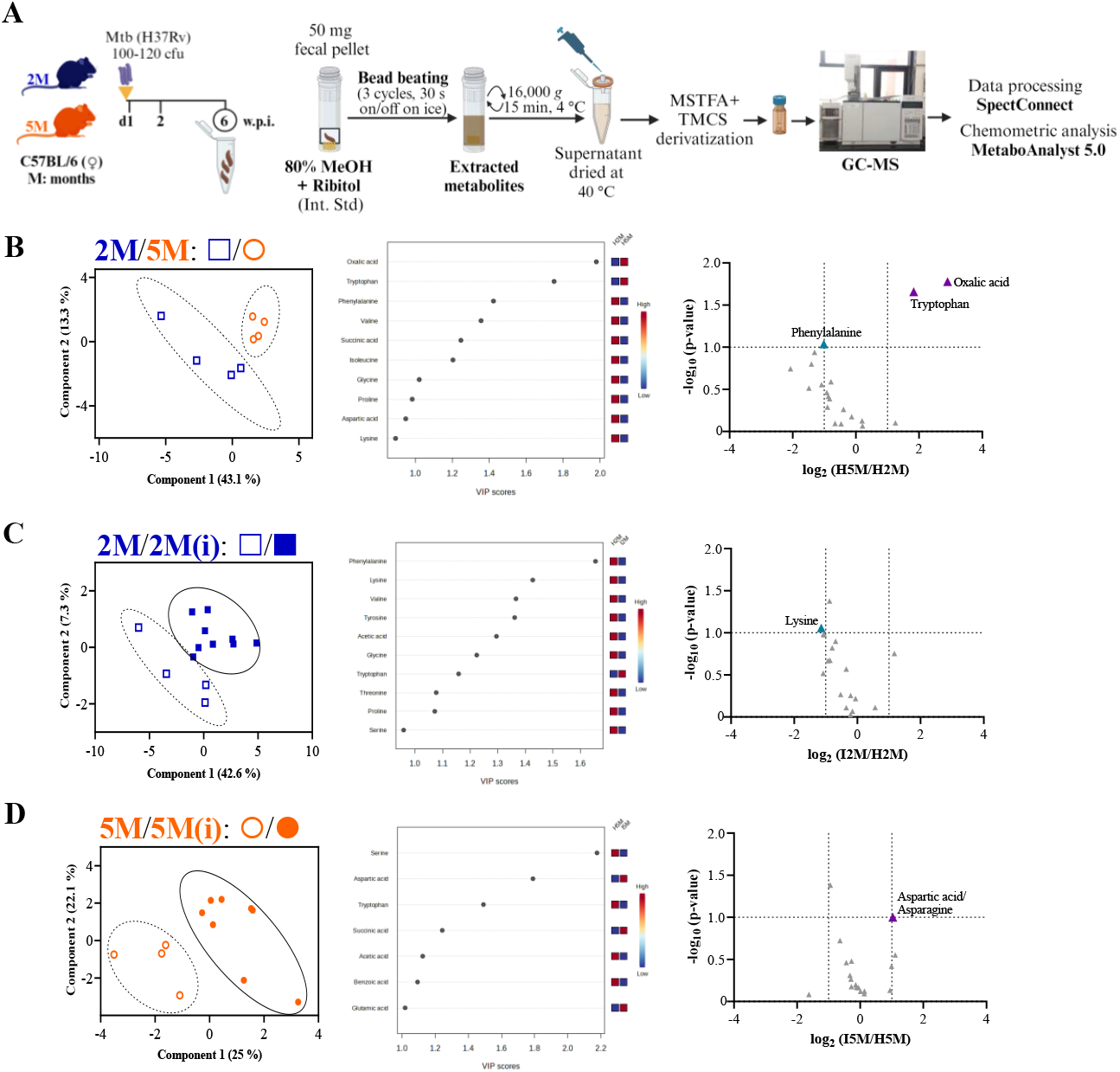
Fecal metabolomics of Mtb infected C57BL/6 mice showed age-specific differences. **A**. Schematic representation for fecal metabolite profiling: Extraction of fecal metabolites, processing of metabolites for Gas Chromatography-mass spectrometry (GC-MS). **B**. Partial least squares-discriminant analysis (PLS-DA) plot representing fecal-specific metabolic differences in healthy (H, n=4/age group) C57BL/6 mice of two age groups, along with analytes having VIP>1 and volcano plot representing deregulated analytes (log_2_FC > ±1 and p-value < 0.1). **C**. PLS-DA plot representing fecal-specific metabolic differences in healthy (H, n=4) and Mtb infected (I, n=9) 2M C57BL/6 mice, along with analytes having VIP>1 and volcano plot. **D**. PLS-DA plot representing fecal-specific metabolic differences in healthy (H, n=4) and Mtb infected (I, n=8) 5M C57BL/6 mice, along with analytes having VIP>1 and volcano plot. M: months; 2M in blue and 5M in orange.

## Discussion

Most pathogens exploit host biochemical machinery and use those products for their basic survival, proliferation, and evading host immune defense. After entry, the tissue-resident immune cells phagocytose the pathogens and expose them to a varied chemical milieu.^6^ In the case of TB, the causative organism, i.e. Mtb, enters the host system through aerosol and gets lodged in the lungs. Initially, alveolar macrophages and tissue-resident type II epithelial cells engulf Mtb, and later, these Mtb-infected cells release chemical messengers attracting the macrophages to the affected organs, like the lungs of Mtb-infected mice and in TB patients. Further, Mtb infection induces tissue inflammation leading to host modulation of its immune system for countering pathogen attack. Perturbed immunometabolism contributed by the local inflammation and chemo-static movement of the immune cells leads to an altered infection-associated molecular profile.^7^ However, this pathogenic attack on the primary tissue might also impact the function and molecular composition of other tissues as well. Few studies have highlighted the metabolic alterations of other non-primary host tissues affected by Mtb infection. In this study, metabolic perturbation in multiple tissues of Mtb aerosol-infected C57BL/6 mice of two age groups was attempted to decipher age-associated host responses. These study groups included commonly used 2M mice belonging to the younger phase of mouse lifespan, which most of the studies use, and another 5M aged mice representing the matured adult phase. Biological ageing significantly impacts metabolic activities, leading to cellular senescence, which is also reported to influence pathological consequences.^8^ Previous report demonstrated an early onset of age-associated metabolic alterations in the lungs and liver of female C67BL/6NRj mice at 6 months of age.^9^

Tissues have their own metabotypes i.e. metabolic phenotype, and understanding pathophysiological changes in different kinds of infection is critical. A tissue-specific metabolome in healthy 2M C57BL/6 mice observed in this study corroborates with the findings in the mouse multiple tissue metabolome database (MMMDB) of healthy 2M C57BL/6J mice.^10^ Metabolite profiling in multiple animal and human tissues has been described by using ultraperformance liquid chromatography (UPLC), ^1^H nuclear magnetic resonance (NMR) and liquid chromatography–high-resolution mass spectrometry (LC– HRMS).^11-13^ Lungs of rats with sepsis analyzed using complementary analytical techniques (LC–MS, GC/MS, and CE–MS) showed deregulated metabolic pathways.^14^

Bacterial infections also accelerate cellular senescence and diminish protective functions, leading to tissue damage impacting other tissue functions.^15^ Metabolic profiling of the biofluids like sputum, serum and urine of TB patients and controls yielded markers explaining TB onset and progression and perturbed lung metabolic phenotype reported in Mtb infected mice models.^16-18^ Therefore, we focused on the lung, i.e., the primary tissue infected with Mtb and others, including the liver, being a highly active metabolic centre of the body and thigh muscle, to understand cachexia-related outcomes while taking the brain as a control. Additionally, the fecal metabolome was also studied for a functional readout of the changes in the gut with Mtb infection.

Upon a low aerosol dose of Mtb infection, the mycobacterial load in the lungs of both 2M and 5M old female C57BL/6 mice was found to be similar till 6 w.p.i. supporting earlier findings on 2M and 6M C57BL/6 mice at 2 w.p.i..^19^ In this study, we observed perturbed metabolism at multi-tissue level with healthy ageing and upon Mtb infection (**Figure 4**). An inverse relationship of aspartic acid/asparagine in the lung (higher) and liver (lower) of healthy 5M mice was observed. Aspartate is reported to increase linearly with age in human lung parenchymal elastin.^20^ It is also known that Mtb exploits asparagine for nitrogen assimilation and resisting acid stress during infection.^21^ Increased lung malonic acid in the Mtb-infected 5M mice corroborates earlier reports at 4 weeks post Mtb infection and is known to inhibit succinate dehydrogenase, leading to decreased cellular respiration.^17,22^ Temporal changes in the lung metabolome of Mtb H37Rv infected ∼2M C57BL/6 mice with higher amino acids and oligopeptides and decreased triacylglycerol levels were reported at 4 and 9 w.p.i..^4^ Decreased mannose levels in the lungs of Mtb-infected 5M mice suggests that the matured age group controls infection efficiently since mannose is required for mycobacterial growth.^23^ Phosphoric acid was elevated in the lungs of Mtb-infected 5M mice, suggesting an altered glucose and protein metabolism in this group. Increased liver xylose in 2M mice suggests increased adipogenesis and improved lipid oxidation upon Mtb infection.^24^ The decreased abundance of liver lysine observed in healthy 5M and Mtb-infected 2M mice suggests these groups control Mtb effectively since lysine regulates the mammalian target of the rapamycin (mTOR) pathway and negatively affects autophagy.^25^ Lysine was also reported to decrease in 5M C57BL6J mice with adenine-induced chronic kidney disease.^26^ On the contrary, higher lung lysine levels were reported in aged (96–104 weeks) wild-type mice with a mixed genetic background of 129/J and C57BL/6J.^27^

**Figure 4.**
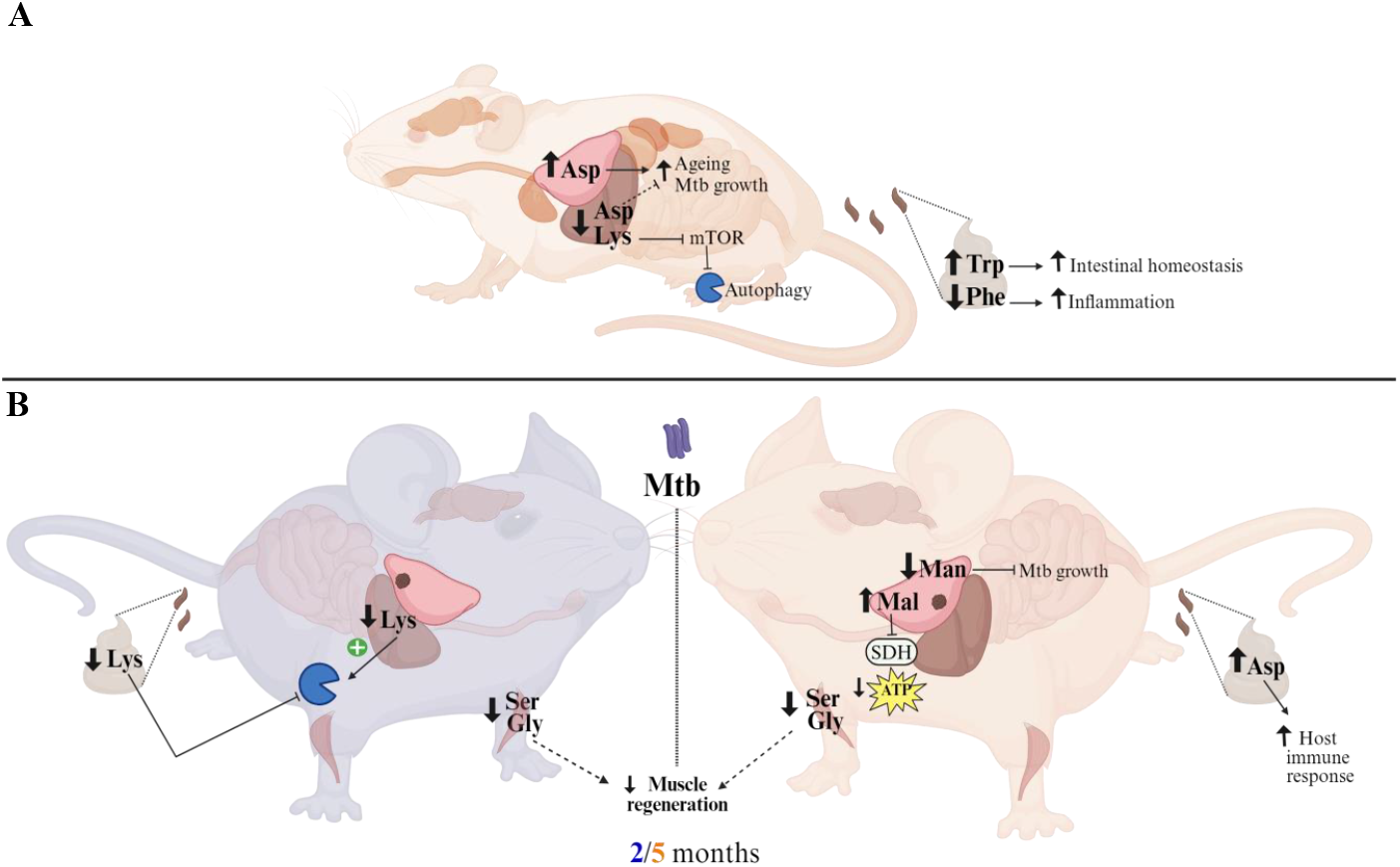
Perturbed tissue and fecal metabolome of C57BL/6 mice upon ageing and upon Mtb infection. Summary of few of the observed changes in tissue (lung, liver, thigh muscle) and feces metabolites in (**A**) healthy 5 months compared to 2 months C57BL/6 mice and (**B**) Mtb H37Rv infected 2 and 5 months compared to respective healthy controls. 2 months in blue and 5 months in orange; Asp: aspartic acid/asparagine, Lys: lysine, mTOR: Mammalian target of rapamycin, Trp: tryptophan, Phe: phenylalanine, Ser: serine, Gly: glycine, Mal: malonic acid, Man: mannose, SDH: succinate dehydrogenase, ATP: adenosine triphosphate. *Figure created with Biorender*

Mtb-infected mice groups showed higher malic acid in the thigh muscle, and it is reported that dietary malic acid supplementation increases the antioxidant capacity and oxidative fibre percentage of weaned piglets, improving meat quality.^28^ Higher thigh muscle oxalic acid observed in Mtb-infected groups corroborates previous reports showing higher pro-inflammatory responses by oxalic acid with a lower glycine: oxalate ratio in atherosclerotic mice.^29^ Mtb-infected mice showed decreased thigh muscle glycine and serine, both of which are known to impair skeletal muscle regeneration in old (>20M) C57BL/6J mice.^30^ While, glycine also demonstrated a protective effect on skeletal muscle in cancer cachexia mouse model and was downregulated in the serum of TB patients.^31,32^ Observed lower glycerol levels in thigh muscle of Mtb-infected mice is a consequence of infection, as higher interstitial glycerol levels are reported in human skeletal muscle.^33^ We did not observe many alterations in the brain metabolic profile post-Mtb infection, possibly due to the method of choice and its rich lipid components.

Age impacts intestinal microbiome composition, which might directly influence the immune system in healthy and disease conditions.^34^ Like microbial diversity, the fecal metabolome provides a functional readout of its activities to mediate a healthy host-microbiome interaction.^35^ Higher fecal oxalic acid levels observed in healthy 5M mice suggest a depletion of oxalic acid degraders like *Oxalobacter formigenes* and corroborates its presence in younger groups.^36,37^ Higher fecal tryptophan metabolites in 5M mice suggest an active immune system since tryptophan contributes to intestinal and systemic homeostasis.^38^ Altered amino acid levels induce intestinal inflammation and are reported to enhance the intestinal mucosal barrier and inhibit oxidative stress by the activation of NF-kB and the production of pro-inflammatory cytokines.^39^ Essential amino acids like phenylalanine and valine, through various pathways, lower the levels of pro-inflammatory cytokines.^39^ We observed lower levels of phenylalanine in the fecal metabolome profile of healthy 5M mice, which suggests a higher inflammation in 5M mice groups. Upon Mtb infection, increased fecal aspartic acid/asparagine levels in 5M mice suggest the unavailability of these nitrogen sources to Mtb, which confers a protective host immune response.^40^ Alternatively, decreased fecal lysine levels in Mtb-infected 2M mice suggest an increased absorption and possibly inhibiting autophagy.

A few limitations of the study include using a smaller metabolite fraction for data analysis to capture amino acids-associated changes, which could be expanded for other metabolic features. Treatment-induced metabolic changes in these mice groups were not attempted and could further shed light on metabolic consequences post TB treatment.

Overall, these findings demonstrated that Mtb infection can have selective metabolic consequences beyond the primary affected organ, i.e. lungs, and further research is needed to fully understand the mechanisms underlying these metabolic changes and their clinical implications.

## Acknowledgements

FP was supported with Junior and Senior Research fellowship from the Department of Biotechnology, Government of India. SC is a Shyama Prasad Mukherjee Fellow supported by the Council of Scientific and Industrial Research, Government of India. The core support from the International Centre for Genetic Engineering and Biotechnology (ICGEB), New Delhi, to RKN, is highly acknowledged. Hossain Md. Faruquee and Sravya Mothe are acknowledged for their help and support during the experiments. Help from the staff of the bio-experimentation facility and Tuberculosis Aerosol Challenge Facility (TACF) at ICGEB; New Delhi, is acknowledged. TACF is supported by the Department of Biotechnology, Government of India. Part of these research findings were presented in Keystone Symposia on Molecular Basis of Healthy Aging, 26-29 March 2023, Breckenridge, CO, United States, World Conference on Lung Health 2022, Virtual, 8-11 November 2022 organized by The Union, 2^nd^ biennial meeting on Sex Differences in the Immune Systems conference, Virtual, 30 November-2 December 2022 organized by Trinity College Dublin & Whitehead Institute and EMBO | EMBL Symposium: Multiomics to Mechanisms: Challenges in Data Integration, 15 – 17 September 2021, Virtual. Some figures have been created with Biorender.com.

## Author Contributions

FP and RKN conceptualized the project. FP and SC carried out animal experiments. FP carried out the tissue metabolomics experiment and data analysis. AD carried out the fecal metabolomics experiment and data analysis. FP and RKN analysed the data, and wrote the original draft, and revised it with the suggestions received from all the co-authors. RKN acquired funding, shared resources, and administrated the project. The manuscript was reviewed and approved by all co-authors.

## Supporting Information

The article contains supplemental information.

## References

1. Snow, K.J., Sismanidis, C., Denholm, J., Sawyer, S.M., & Graham, S.M. (2018). The incidence of tuberculosis among adolescents and young adults: A global estimate. Eur. Respir. J. 51: 1702352.

2. Morabia, A. (2014). Snippets from the past: cohort analysis of disease rates – Another piece in a seemingly still incomplete puzzle. Am. J. Epidemiol. 180: 189–196.

3. Flurkey, K., Currer, J.M., Harrison, D.E., & Fox, J.G. (2007). The mouse in biomedical research. American College of Laboratory Animal Medicine series. Elsevier, AP: Amsterdam, 637–672.

4. Fernandez-Garcia, M., Rey-Stolle, F., Boccard, J., Reddy, V. P., García, A., Cumming, B. M., … & Barbas, C. (2020). Comprehensive examination of the mouse lung metabolome following Mycobacterium tuberculosis infection using a multiplatform mass spectrometry approach. J. Proteome. Res. 19: 2053–2070.

5. Somashekar, B.S., Amin, A.G., Rithner, C.D., Troudt, J., Basaraba, R., Izzo, A., Crick, D.C., & Chatterjee, D. (2011). Metabolic profiling of lung granuloma in Mycobacterium tuberculosis infected guinea pigs: ex vivo 1H magic angle spinning NMR studies. J. Proteome Res. 10: 4186–4195.

6. Russell, D.G., VanderVen, B.C., Lee, W., Abramovitch, R.B., Kim, M.J., Homolka, S., … & Rohde, K.H. (2010). Mycobacterium tuberculosis wears what it eats. Cell Host Microbe 8: 68–76.

7. Cui, L., Zheng, D., Lee, Y. H., Chan, T. K., Kumar, Y., Ho, W.E., … & Ong, C. N. (2016). Metabolomics investigation reveals metabolite mediators associated with acute lung injury and repair in a murine model of influenza pneumonia. Sci. Rep. 6: 26076.

8. Campisi, J. (2013). Aging, cellular senescence, and cancer. Annu. Rev. Physiol. 75: 685–705.

9. Bresilla, D., Habisch, H., Pritišanac, I., Zarse, K., Parichatikanond, W., Ristow, M., Madl, T., & Madreiter-Sokolowski, C.T. (2022). The sex-specific metabolic signature of C57BL/6NRj mice during aging. Sci. Rep. 12: 21050.

10. Sugimoto, M., Ikeda, S., Niigata, K., Tomita, M., Sato, H., & Soga, T. MMMDB: Mouse multiple tissue metabolome database. (2012). Nucleic Acids Res. 40(Database issue): D809–D814.

11. Want, E.J., Masson, P., Michopoulos, F., Wilson, I.D., Theodoridis, G., Plumb, R.S., Shockcor, J., Loftus, N., Holmes, E., & Nicholson, J.K. (2013). Global metabolic profiling of animal and human tissues via UPLC-MS. Nat. Protoc. 8: 17–32.

12. Beckonert, O. et al. (2007). Metabolic profiling, metabolomic and metabonomic procedures for NMR spectroscopy of urine, plasma, serum and tissue extracts. Nat. Protoc. 2: 2692–2703.

13. Burlikowska, K., Stryjak, I., Bogusiewicz, J., Kupcewicz, B., Jaroch, K., & Bojko, B. (2020). Comparison of metabolomic profiles of organs in mice of different strains based on SPME-LC-HRMS. Metabolites 10: 255.

14. Naz, S., Antonia G., & Coral, B. (2013). Multiplatform analytical methodology for metabolic fingerprinting of lung tissue. Anal. Chem. 85: 10941–10948.

15. Teissier, T., Boulanger, E., & Cox, L. S. (2022). Interconnections between inflammageing and immunosenescence during ageing. Cells 11: 359.

16. du--Preezm I., Luies, L., & Loots, D.T. (2019). The application of metabolomics toward pulmonary tuberculosis research. Tuberculosis 115: 126–139.

17. Fernández-García, M., Rojo, D., Rey-Stolle, F., García, A., & Barbas, C. (2018). Metabolomic-based methods in diagnosis and monitoring infection progression. In Metabolic Interaction in Infection, 1st ed.; Silvestre R..; Torrado E., Eds..; Springer: 283–315.

18. Mirsaeidi, M., Banoei, M.M., Winston, B.W., & Schraufnagel, D.E. (2015). Metabolomics: Applications and promise in mycobacterial disease. Ann. Am. Thorac. Soc. 212: 1278–1287.

19. Vesosky, B., Flaherty, D. K., Rottinghaus, E. K., Beamer, G. L., & Turner, J. (2006). Age dependent increase in early resistance of mice to Mycobacterium tuberculosis is associated with an increase in CD8 T cells that are capable of antigen independent IFN-γ production. Exp. Gerontol. 41: 1185–1194.

20. Shapiro, S.D., Endicott, S.K., Province, M.A., Pierce, J.A., & Campbell, E.J. (1991). Marked longevity of human lung parenchymal elastic fibers deduced from prevalence of D-aspartate and nuclear weapons-related radiocarbon. J. Clin. Invest. 87: 1828–1834.

21. Gouzy, A., Larrouy-Maumus, G., Bottai, D., Levillain, F., Dumas, A., Wallach, J.B., Caire-Brandli, I., de Chastellier, C., Wu, T.D., Poincloux, R., & Brosch, R. (2014). Mycobacterium tuberculosis exploits asparagine to assimilate nitrogen and resist acid stress during infection. PLoS Pathog. 10: p.e1003928.

22. Pardee, A.B., & Potter, V.R. (1949). Malonate inhibition of oxidations in the Krebs tricarboxylic acid cycle. J. Biol. Chem. 178: 241–250.

23. Patterson, J.H., Waller, R.F., Jeevarajah, D., Billman-Jacobe, H., & McConville, M.J. (2003). Mannose metabolism is required for mycobacterial growth. Biochem. J. 372: 77–86.

24. Lim, E., Lim, J.Y., Shin, J.H., Seok, P.R., Jung, S., Yoo, S.H., & Kim, Y. (2015). d-Xylose suppresses adipogenesis and regulates lipid metabolism genes in high-fat diet–induced obese mice. Nutr. Res. 35: 626–636.

25. Schroeder, S. et al. (2014). Metabolites in aging and autophagy. Microb. Cell 1: 110–114.

26. Khattri, R.B., Thome, T., & Ryan, T.E. (2021). Tissue-specific 1H-NMR metabolomic profiling in mice with adenine-induced chronic kidney disease. Metabolites 11: 45.

27. Zhang, F., Kerbl-Knapp, J., Akhmetshina, A., Korbelius, M., Kuentzel, K.B., Vujić, N., Hörl, G., Paar, M., Kratky, D., Steyrer, E., & Madl, T. (2021). Tissue-specific landscape of metabolic dysregulation during ageing. Biomolecules 11: 235.

28. Zhang, X., Chen, M., Yan, E., Wang, Y., Ma, C., Zhang, P., & Yin, J. (2022). Dietary malic acid supplementation induces skeletal muscle fiber-type transition of weaned piglets and further improves meat quality of finishing pigs. Front. Nutr. 8: 825495.

29. Liu, Y., Zhao, Y., Shukha, Y., Lu, H., Wang, L., Liu, Z., Liu, C., Zhao, Y., Wang, H., Zhao, G., & Liang, W. (2021). Dysregulated oxalate metabolism is a driver and therapeutic target in atherosclerosis. Cell Rep. 36: 109420.

30. Gheller, B.J., Blum, J.E., Lim, E.W., Handzlik, M.K., Fong, E.H.H., Ko, A.C., Khanna, S., Gheller, M.E., Bender, E.L., Alexander, M.S., & Stover, P.J. (2021). Extracellular serine and glycine are required for mouse and human skeletal muscle stem and progenitor cell function. Mol. Metab. 43: 101106.

31. Zhou, A., Ni, J., Xu, Z., Wang, Y., Lu, S., Sha, W., Karakousis, P.C., & Yao, Y.F. (2013). Application of 1H NMR spectroscopy-based metabolomics to sera of tuberculosis patients. J. Proteome Res. 12: 4642–4649.

32. Ham, D.J., Murphy, K.T., Chee, A., Lynch, G.S., & Koopman, R. (2014). Glycine administration attenuates skeletal muscle wasting in a mouse model of cancer cachexia. Clin. Nutr. 33: 448–458.

33. Maggs, D.G., Jacob, R., Rife, F., Lange, R., Leone, P., During, M.J., Tamborlane, W.V., & Sherwin, R.S. (1995). Interstitial fluid concentrations of glycerol, glucose, and amino acids in human quadricep muscle and adipose tissue. Evidence for significant lipolysis in skeletal muscle. J. Clin. Invest. 96: 370–377.

34. McHugh, D., & Gil, J. (2018). Senescence and aging: Causes, consequences, and therapeutic avenues. J. Cell Biol. 217: 65–77.

35. Zierer, J., Jackson, M.A., Kastenmüller, G., Mangino, M., Long, T., Telenti, A., Mohney, R.P., Small, K.S., Bell, J.T., Steves, C.J., & Valdes, A.M. (2018). The fecal metabolome as a functional readout of the gut microbiome. Nat. Genet. 50: 790–795.

36. Li, X., Ellis, M. L., & Knight, J. (2015). Oxalobacter formigenes colonization and oxalate dynamics in a mouse model. Appl. Environ. Microbiol. 81: 5048–5054.

37. Barnett, C., Nazzal, L., Goldfarb, D. S., & Blaser, M. J. (2016). The presence of Oxalobacter formigenes in the microbiome of healthy young adults. J. Urol. 195: 499–506.

38. Roager, H. M., & Licht, T. R. (2018). Microbial tryptophan catabolites in health and disease. Nat. Comm. 9: 3294.

39. He, F., Wu, C., Li, P., Li, N., Zhang, D., Zhu, Q., Ren, W., & Peng, Y. (2018). Functions and signaling pathways of amino acids in intestinal inflammation. Biomed Res. Int. 2018.

40. Borah, K., Beyß, M., Theorell, A., Wu, H., Basu, P., Mendum, T.A., Nӧh, K., Beste, D.J., & McFadden, J. (2019). Intracellular Mycobacterium tuberculosis exploits multiple host nitrogen sources during growth in human macrophages. Cell Rep. 29: 3580–3591.

