## Supplementary Figs S1 to S7, Supplementary Table S1 for "Lung *Mycobacterium tuberculosis* infection perturbs metabolic pathways in non-pulmonary tissues"

<u>**Supplemental Information**</u>

**A**

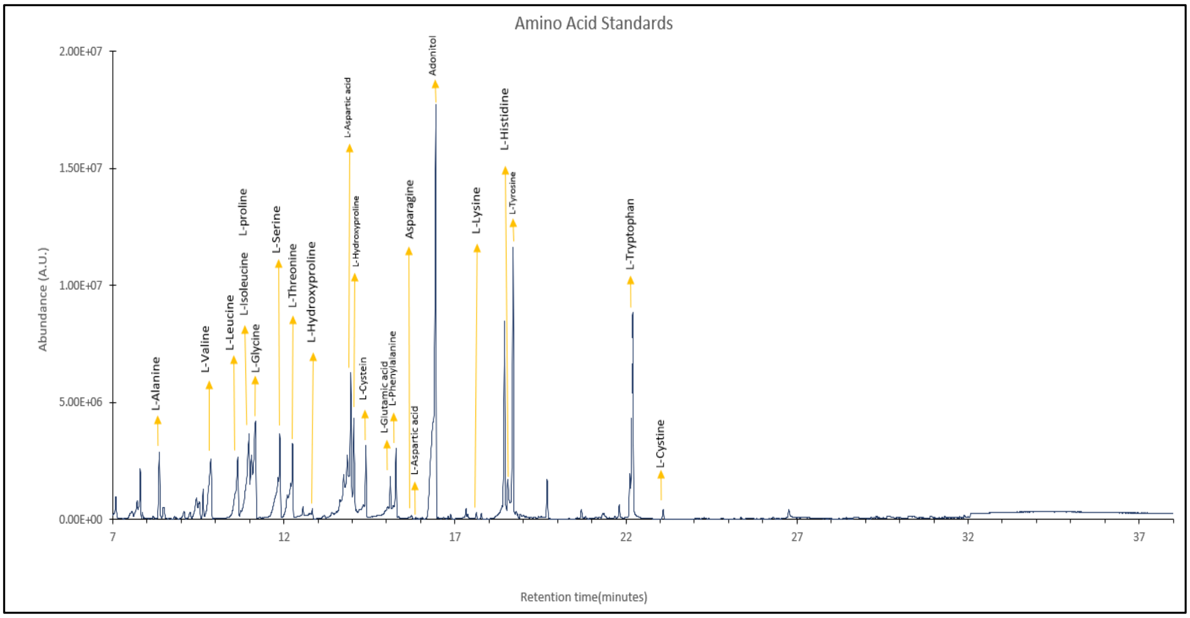


**B**

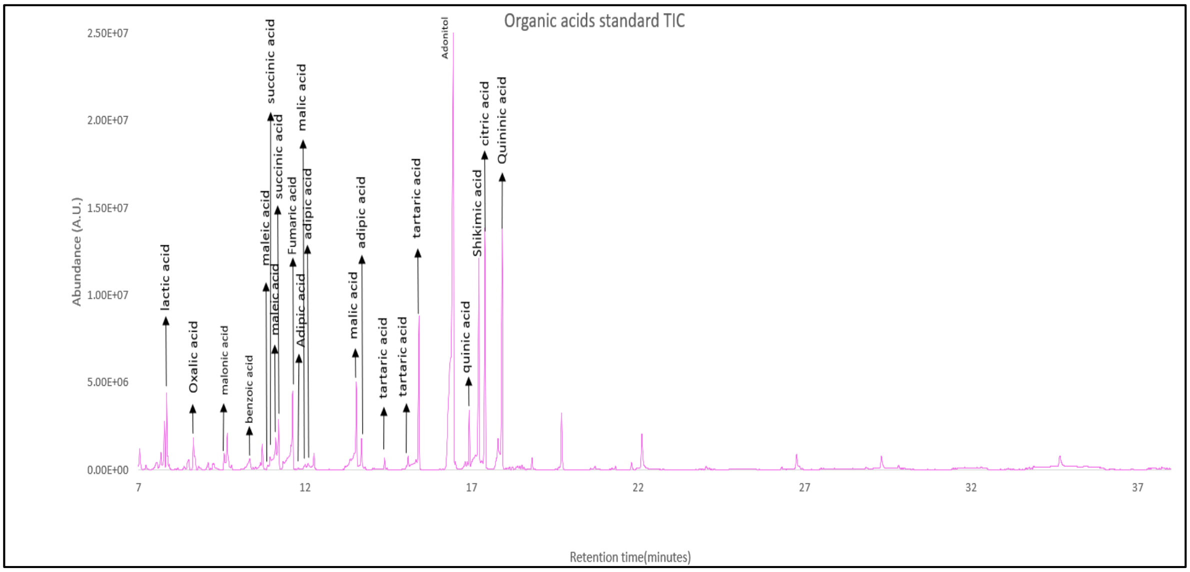


**Supplementary Figure S1**. **Commercial standards used in fecal metabolomics study**. Total ion chromatogram (TIC) of available (**A**) amino acids and (**B**) organic acids.

A

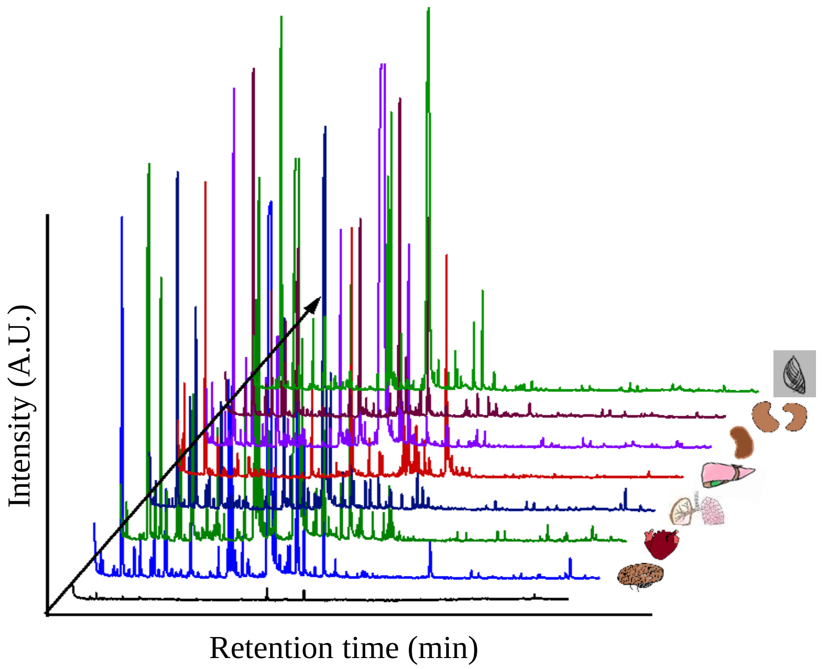


B

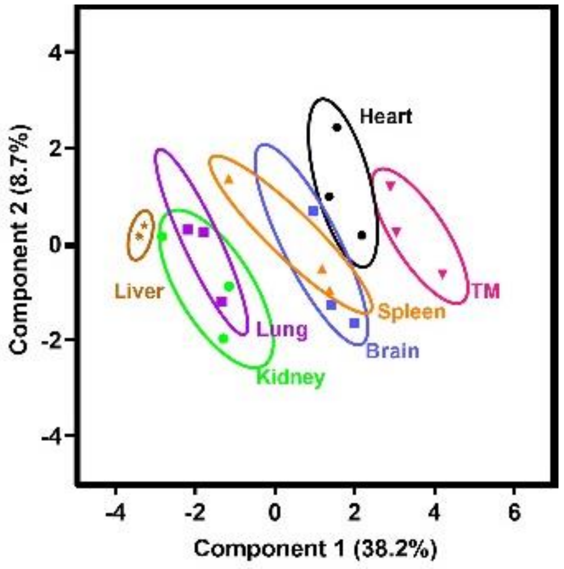


**Supplementary Figure S2**. **Optimization of tissue metabolomics study in healthy C57BL/6 mice**. **A**. Representative total ion chromatogram (TIC) of extracted tissue metabolites harvested from healthy 2 months female C57BL/7 mice. **B**. Partial least squares-discriminant analysis (PLS-DA) plot representing tissue-specific metabolic differences (Ovals used are only for presenting tissue clusters without considering statistical confidence intervals); n= 3/tissue.

**A**

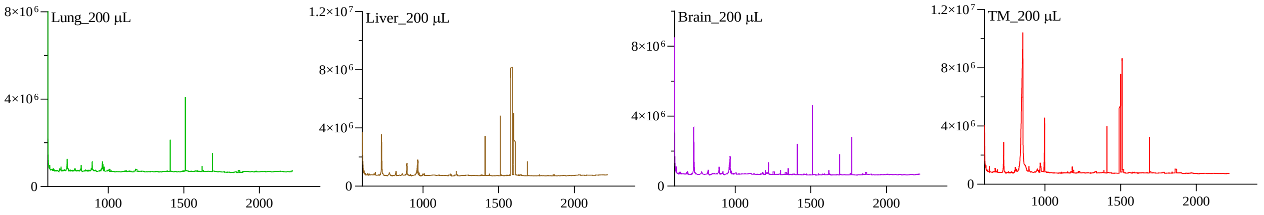


**B**

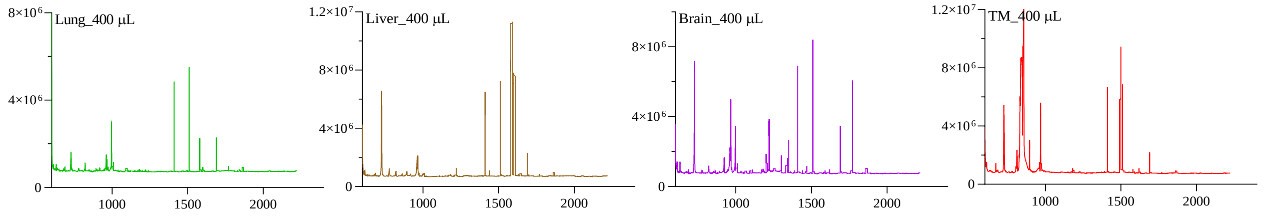


**C**

| Tissue | Volume (µL) | Features |
|---|---|---|
| Lung | 200 | 69 |
| | 400 | 86 |
| Liver | 200 | 79 |
| | 400 | 104 |
| Brain | 200 | 71 |
| | 400 | 119 |
| TM | 200 | 86 |
| | 400 | 95 |

**Supplementary Figure S3. Volume optimization in tissue metabolomics study**.

Total ion chromatogram (TIC) of the extracted metabolites from lung, liver, brain and thigh muscle (**A**) from 200 µL and (**B**) 400 µL. **C**. Number of features identified from 200 µL and 400 µL of the extracted tissue metabolites.

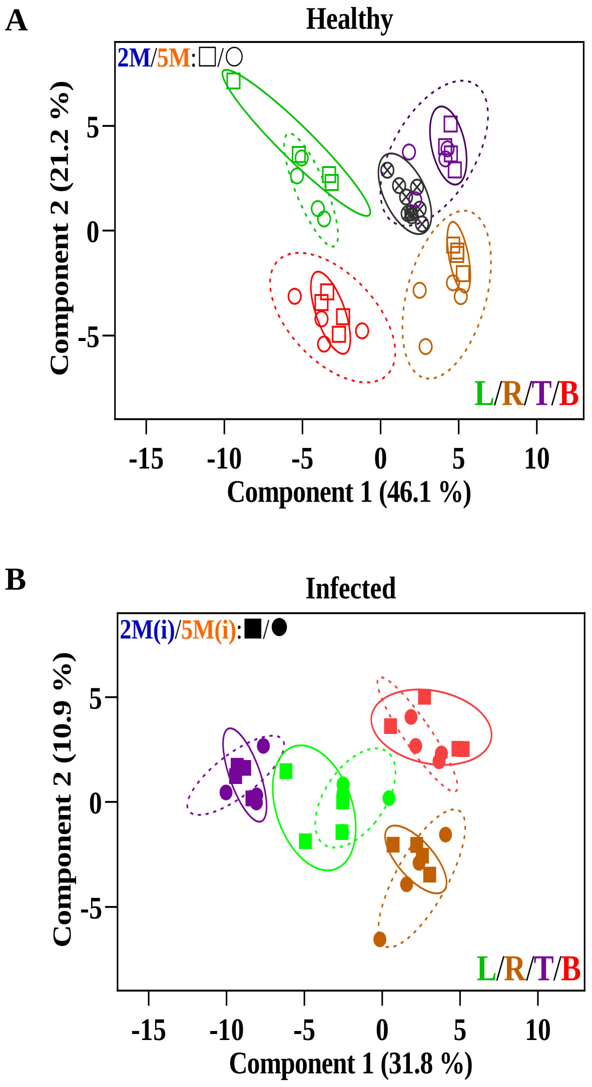


**Supplementary Figure S4**. **Tissue metabolomics of Mtb infected C57BL/6 mice showed age-specific differences**. **A**. Partial least squares-discriminant analysis (PLS-DA) plot (n= 4/age group) representing tissue-specific metabolic differences belonging to 2M and 5M age groups of C57BL/6 mice; QC samples represented in black circle. **B**. PLS-DA plot (n= 4/age group) representing tissue-specific metabolic differences in Mtb H37Rv infected tissues of 2M and 5M age groups of C57BL/6 mice. M: months.

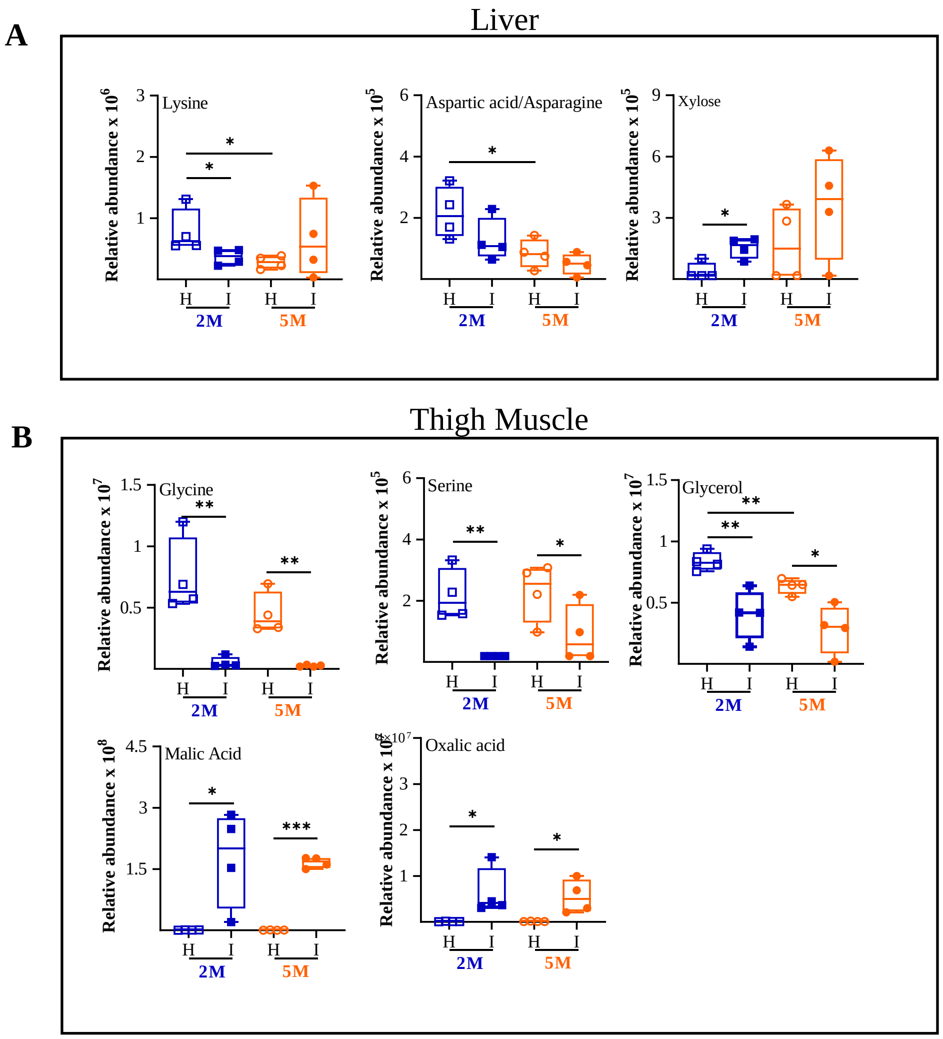


**Supplementary Figure S5**. **Mtb infection perturbs liver and thigh muscle metabolome**. Relative abundance of key metabolites showing changes in (**A**) liver and (**B**) thigh muscle of healthy (H) and Mtb H37Rv infected (I) C57BL/6 mice of two age groups. n=4/age group/condition; M: months; 2M in blue and 5M in orange. p-value: * ≤0.01 , ** ≤0.01 and *** ≤0.001 at 90% confidence interval.

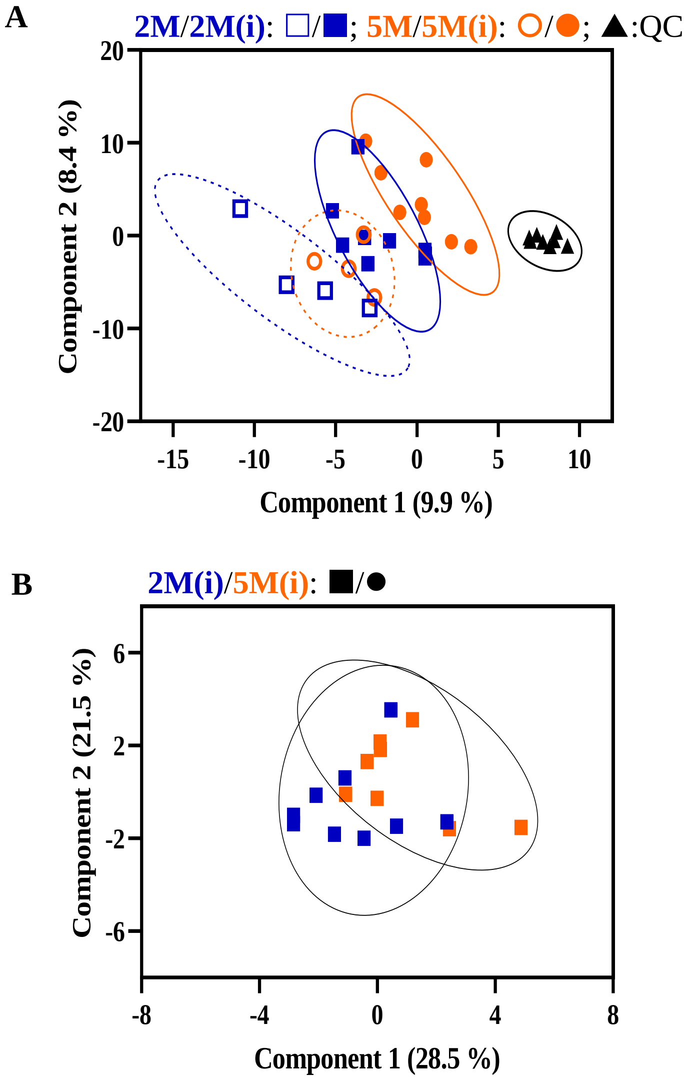


**Supplementary Figure S6**. **Age-specific differences in fecal metabolites of Mtb infected C57BL/6 mice**. **A**. Partial least squares-discriminant analysis (PLS-DA) plot representing tissue-specific metabolic differences belonging to healthy and Mtb H37Rv infected 2M and 5M age groups of C57BL/6 mice; QC samples represented in black triangle. **B**. PLS-DA plot representing tissue-specific metabolic differences in Mtb infected tissues of 2M and 5M age groups of C57BL/6 mice. Healthy (open symbols, n= 4 per age group); Mtb H37Rv infected (closed symbols, n=9 for 2M mice and n=8 for 5M); M: months; 2M in blue and 5M in orange.

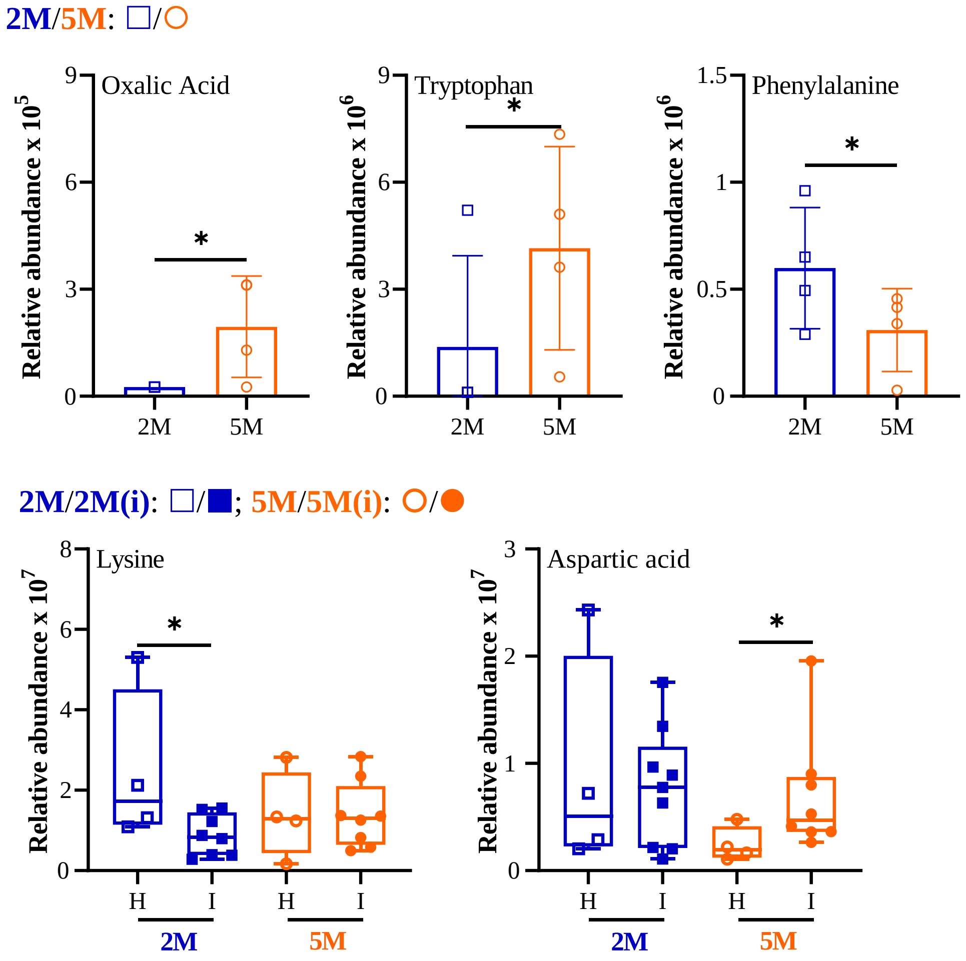


**Supplementary Figure S7**. **Deregulated amino acids observed in the fecal metabolome of 5M C57BL/6 mice**. Histograms showing the relative abundance of oxalic acid, tryptophan, phenylalanine, lysine and aspartic acid in 2M and 5M C57BL/6 mice. Healthy (H, n= 4 per age group); Mtb H37Rv infected (I, n=9 for 2M mice and n=8 for 5M); M: months; 2M in blue and 5M in orange; p-values: * ≤0.1 at 90% confidence interval.

**Supplementary Table S1**. **A**. Total number of variables used for comparative tissue metabolome study in 2M and 5M C57BL/6 mice. **B**. List of identified variables covering amino acid metabolism used for comparative tissue metabolome analysis.

**A**

| **Tissue** | **2M** | **5M** |
|---|---|---|
| Lung | 47 | 57 |
| Liver | 61 | 62 |
| Thigh Muscle | 61 | 58 |
| Brain | 60 | 59 |

**B**

| **S.No.** | **Variable** |
|---|---|
| 1 | 2-Butenedioic acid |
| 2 | 2-Hydroxyisocaproic acid |
| 3 | Allonic acid |
| 4 | Aspartic acid/Asparagine |
| 5 | Butanoic acid |
| 6 | Citrulline |
| 7 | Creatinine |
| 8 | Xylose |
| 9 | Mannose |
| 10 | Erythro-Pentonic acid |
| 11 | Oxalic acid |
| 12 | Glutamic acid/Glutamine |
| 13 | Glycerol |
| 14 | Glycine |
| 15 | Hydroxypyruvic acid |
| 16 | Lysine |
| 17 | Malic acid |
| 18 | Phenylalanine |
| 19 | Pentanoic acid |
| 20 | Phosphoric acid |
| 21 | Proline |
| 22 | Malonic acid |
| 23 | Propanoic acid |
| 24 | Pyroglutamic acid |
| 25 | Serine |
| 26 | Threonine |
| 27 | Trans-9-Octadecenoic acid |
| 28 | Urea |
| 29 | Valine |
